# canvasXpress: A versatile interactive high-resolution scientific multi-panel visualization toolkit

**DOI:** 10.1101/186213

**Authors:** Baohong Zhang, Shanrong Zhao, Isaac Neuhaus

## Abstract

**To the Editor:** CanvasXpress (https://canvasxpress.org) was developed as the core visualization component for bioinformatics and systems biology analysis at Bristol-Myers Squibb and further enhanced by scientists around the world and served as a key visualization engine for many popular bioinformatics tools^1,2,3,4,5,6^. It offers a rich set of interactive plots to display scientific and genomics data, such as oncoprint of cancer mutations, heatmap, 3D scatter, violin, radar, and profile plots (Figure 1, canvasXpress plots arranged by canvasDesigner https://baohongz.github.io/canvasDesigner). Recently, the reproducibility and usability of the package in real world bioinformatics and clinical use cases have been improved significantly witnessed by continuous add-on features and wide adoption of the toolkit in the scientific communities. Furthermore, It is the first noteworthy package harmonizing real time interactive exploring and analyzing of big data, full-fledged customizing of look-n-feel, and producing multi-panel publication-ready figures in PDF format simultaneously.

---

Emphasizing reproducible research, canvasXpress captures all programmatic and user interactions including modifications done through the menus, on plot controls and the comprehensive configurator as illustrated at https://canvasxpress.org/html/user-interface.html. The customization is recorded in real time as the user modifying a graph and is embedded in the saved SVG file as metadata. The file can then be opened in a webpage containing a canvasXpress panel to reproduce the exact plot. Please see a step-by-step example at https://canvasxpress.org/html/reproducible-research.html.

**Figure 1.**
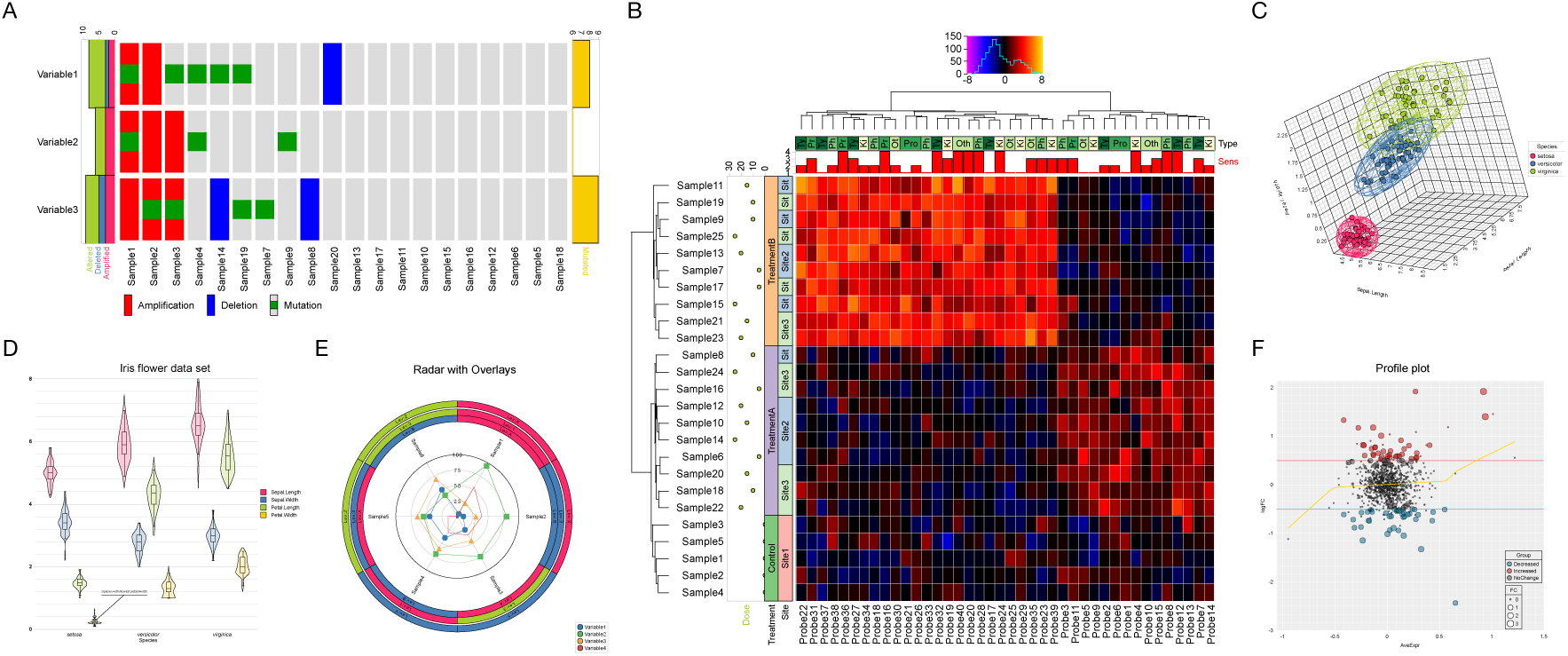
Versatile plots generated by canvasXpress. A) Oncoprint of cancer mutations; B) Gene expression heatmap with comprehensive annotations; C) 3D scatter plot; D) Violin plot; E) Radar plot with annotations; F) Profile plot of gene expression.

CanvasXpress also includes a standalone unobtrusive data table and a filtering widget to allow data exploration and filtering similar to those only seen in high-end commercial tools. Data can be easily sorted, grouped by multiple levels, segregated, transposed, transformed or clustered dynamically. The fully customizable mouse events as well as the zooming, panning and drag-n-drop capabilities are features that make this library capable of further expansion of functionalities.

CanvasXpress is a standalone JavaScript library that works in all modern browsers on mobile, tablets and desktop devices. The following example shows the basic usage which consists of four elements: the JavaScript and the CSS framework in the <head> section; the data also in the <head>; a <canvas> element in the <body>; and lastly including a mechanism to instantiate the canvasXpress object which in this example is a call to a function triggered after loading the web page.

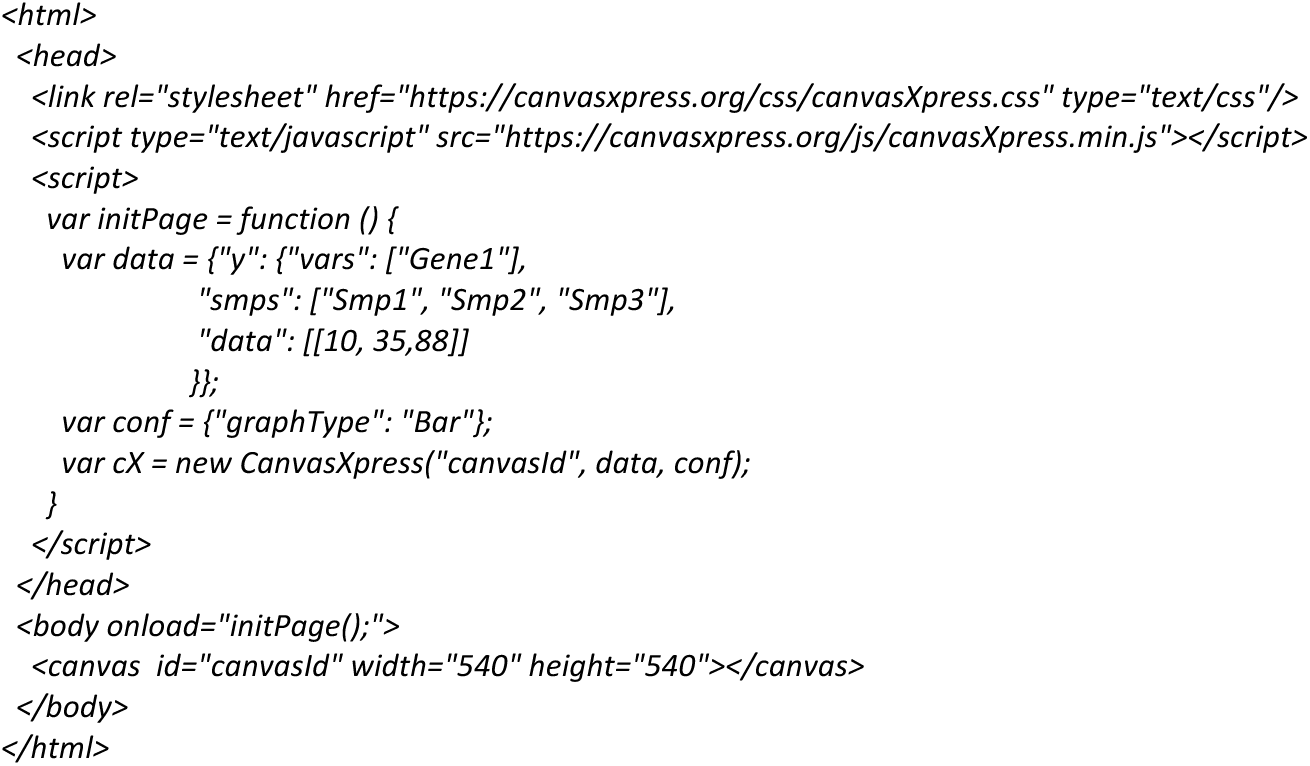

CanvasXpress can be now simply used within R at the console to generate conventional plots, in R-Studio or seamlessly embedded in Shiny web applications. A list of examples of the canvasXpress R library including the mouse events, zooming, and broadcasting capabilities are included under the shiny directory in the github package (https://github.com/neuhausi/canvasXpress). This R library was created with the htmlwidgets package.

Facilitated by canvasDesigner, the bundled graphics layout tool written in JavaScript, the end user can easily arrange any number of different types of plots in SVG format outputted by canvasXpress or other sources on a HTML page. Each individual plot could be adjusted in size and positioned freely. At last, printing as PDF can generate a high-resolution multi-panel plot as shown in Figure 1, which is a standard option on all modern Internet browsers. This design tool relieves scientists from days if not weeks of struggle of using multiple tools, sometime even programming in R or some scripting languages in order to get job done.

CanvasXpress is the first open source package with unprecedented functionalities that are not even seen in expensive commercial tools. It empowers biologists who don’t possess advanced computer skills to perform analysis and generate complex publication-ready figures required by the prestigious journals at ease in the big data era.

## AUTHOR CONTRIBUTIONS

I.N. conceived the project. I.N. wrote the canvasXpress software. B.Z. wrote canvasDesigner tool. B.Z, S.Z. and I.N. wrote the manuscript.

## Supplementary of canvasXpress

### Web URLs to tools and online user guide

Google Chrome (https://www.google.com/chrome) is recommended for optimal use of the web based tools.

canvasXpress: https://canvasxpress.org

canvasXpress source code: https://github.com/neuhausi/canvasXpress

canvasXpress download: https://canvasxpress.org/html/download.html

canvasXpress user interface: https://canvasxpress.org/html/user-interface.html

canvasXpress examples: https://canvasxpress.org/html/bar-1.html

canvasDesigner: https://baohongz.github.io/canvasDesigner

canvasDesigner source code: https://github.com/baohongz/canvasDesigner

canvasDesigner example #1: all SVG files are generated by canvasXpress. https://baohongz.github.io/canvasDesigner/example1.html

canvasDesigner example #2: mixed SVG files from canvasXpress and other sources. https://baohongz.github.io/canvasDesigner/example2.html

Example SVG files: https://github.com/baohongz/canvasDesigner/tree/gh-pages/SVG

Inkscape: Edit SVG files, convert image files in other formats to SVG format. https://inkscape.org

SVGOMG: Optimize SVG files. https://jakearchibald.github.io/svgomg

### SVG (Scalable Vector Graphics) files generated by canvasXpress

All plots generated by canvasXpress can be saved as SVG file by following the menu.

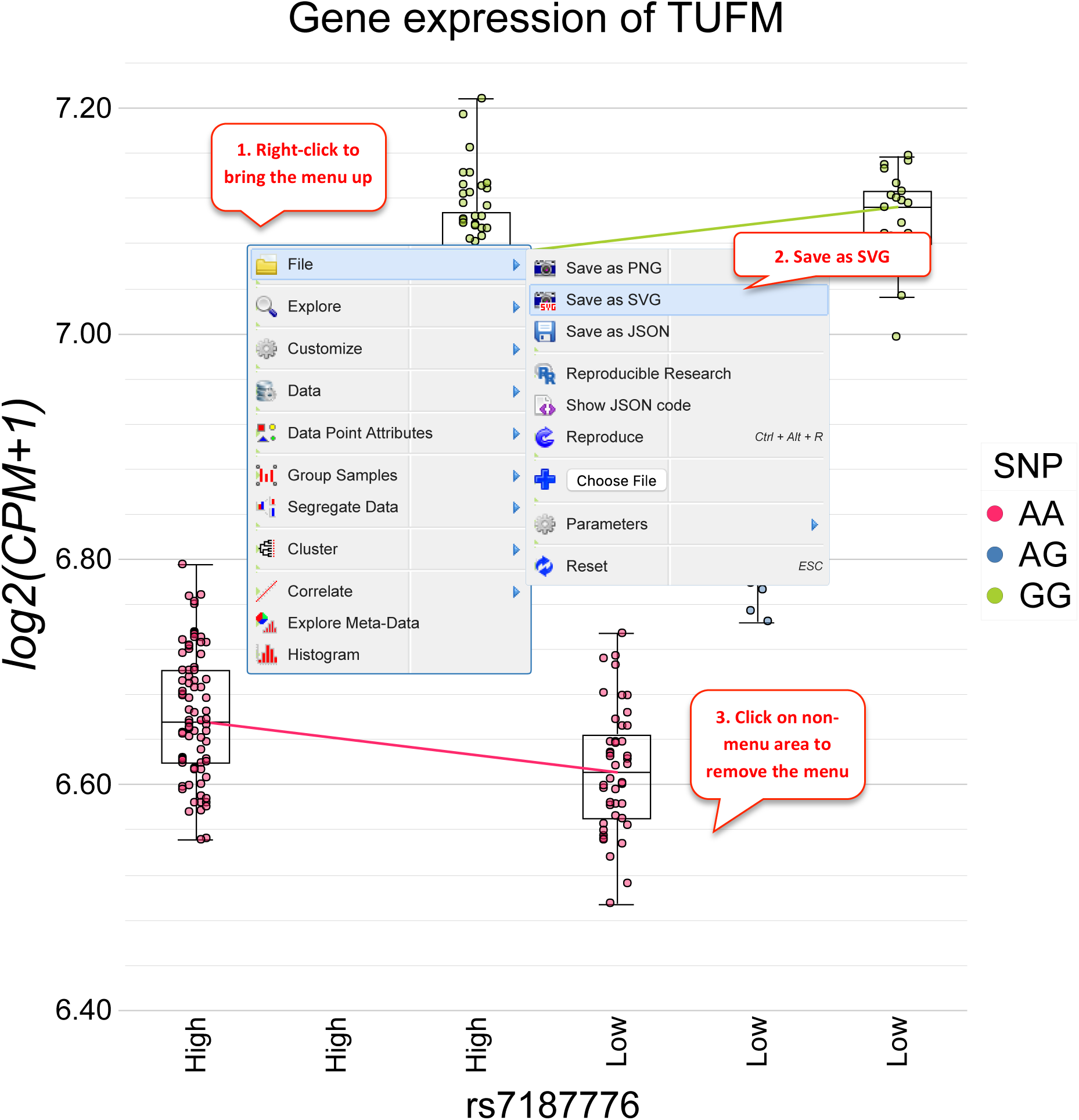

### Layout multiple SVG files by canvasDesigner

The HTML based tool can easily arrange multiple plots in SVG format outputted by canvasXpress or other sources. Each individual plot could be adjusted in size and positioned freely.

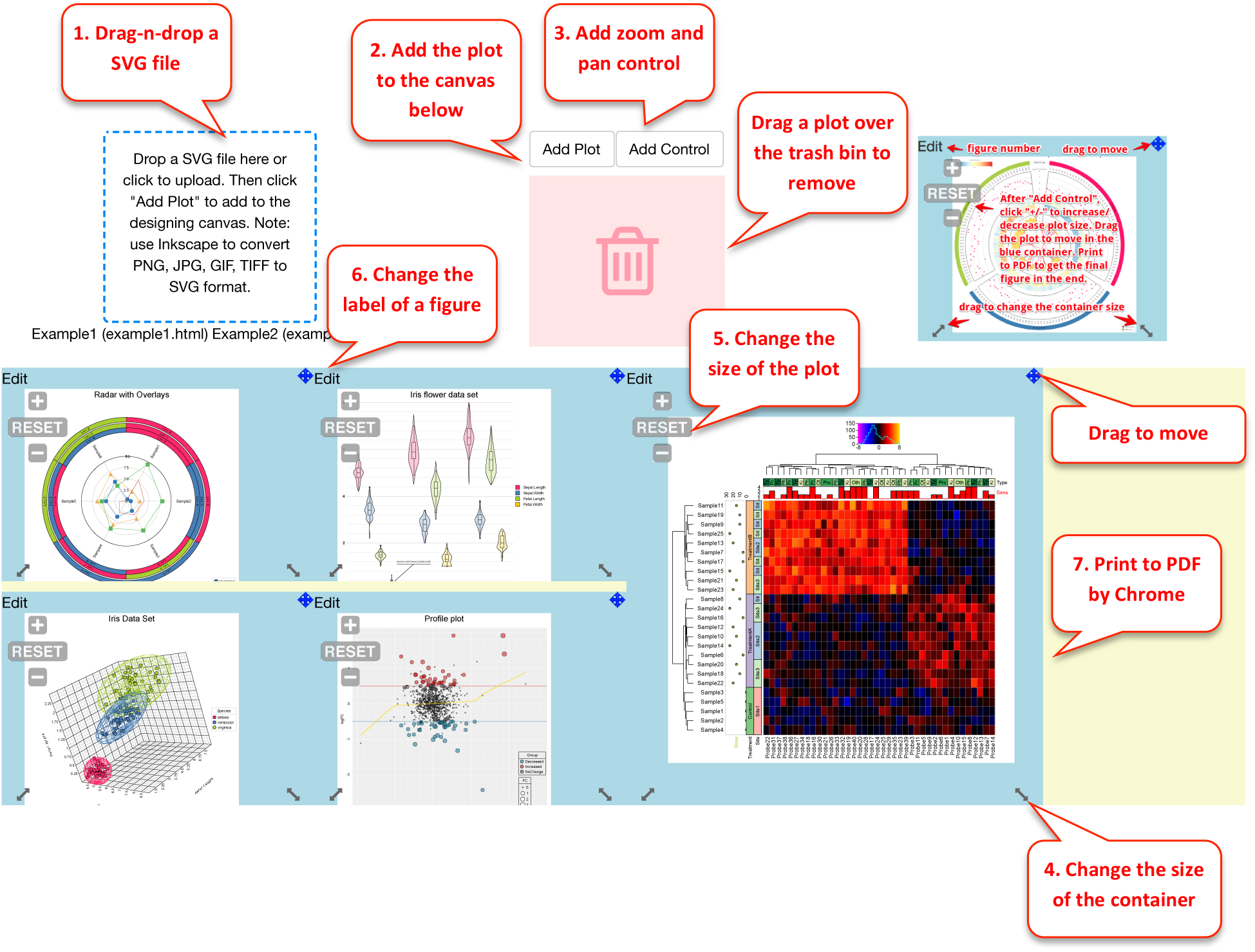

canvasDesigner example #1: all SVG files are generated by canvasXpress. https://baohongz.github.io/canvasDesigner/examplel.html

canvasDesigner example #2: mixed SVG files from canvasXpress and other sources. https://baohongz.github.io/canvasDesigner/example2.html

### Inkscape to annotate and convert images to SVG if needed

Inkscape (https://inkscape.org) is powerful open-source vector graphics editor. You can add text annotations to jpeg, png, gif, and tiff image and save it as “Plain SVG” file. The example is to show how to add annotations to a TIFF image.

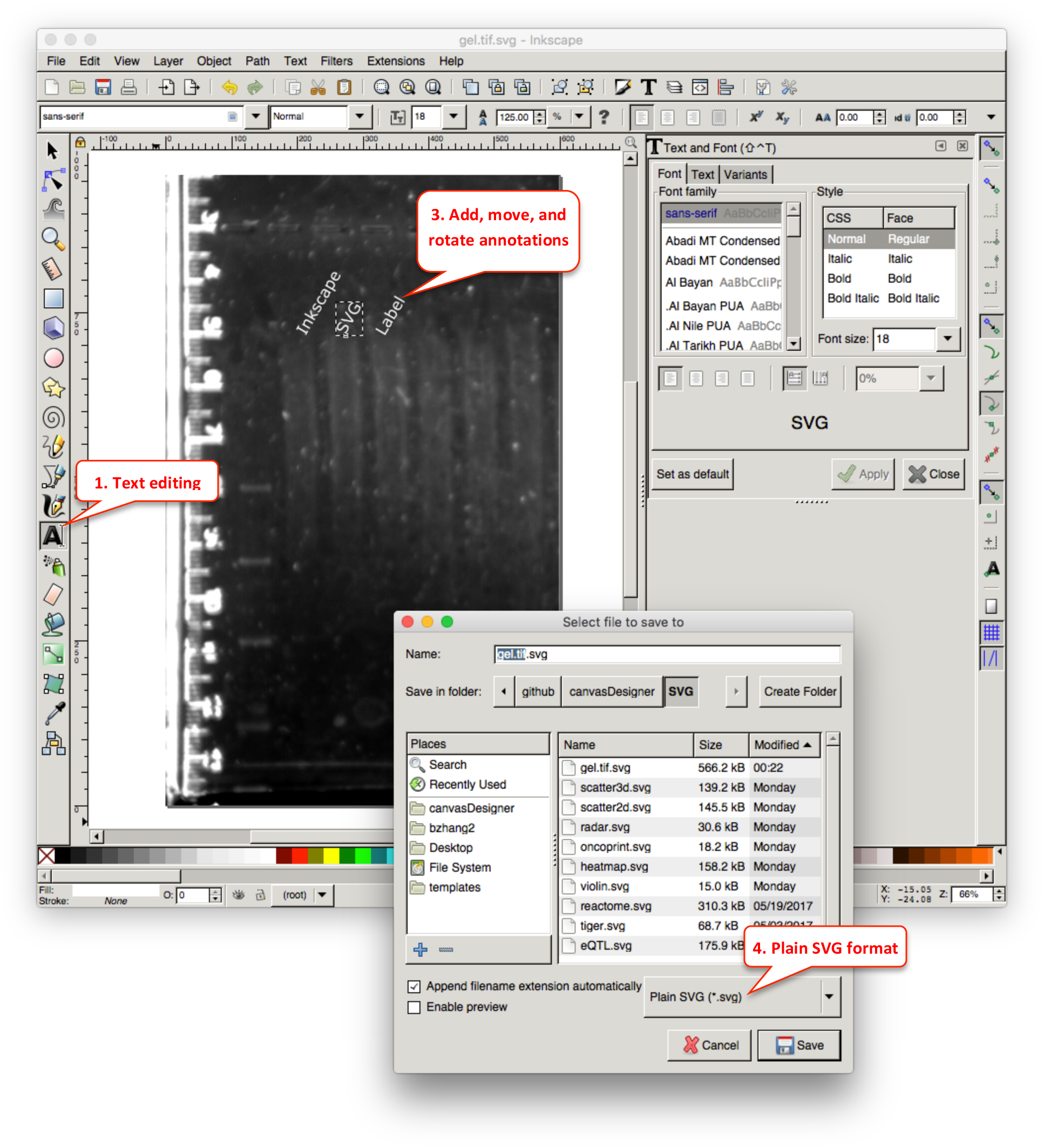

### Optimize SVG files by SVGOMG

SVG files, especially exported from various tools, usually contain a lot of redundant and useless information such as editor metadata, comments, hidden elements, default or non-optimal values and other stuff that can be safely removed or converted by the online tool SVGOMG (https://jakearchibald.github.io/svgomg) without affecting SVG rendering result.

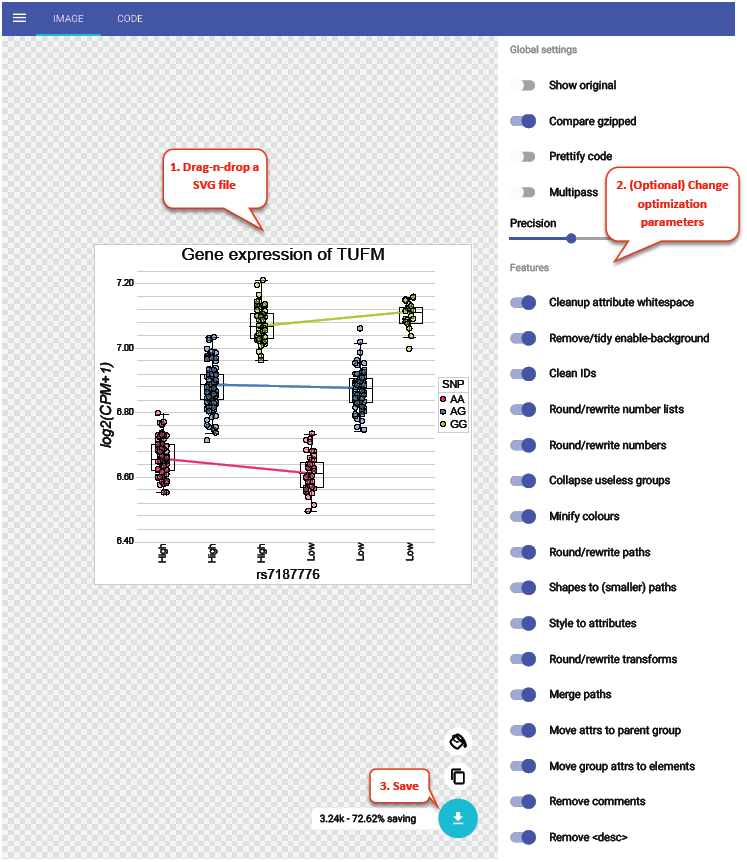

